# Lower limb pointing to assess intersegmental dynamics after incomplete spinal cord injury and the associated role of proprioceptive impairments

**DOI:** 10.1101/2024.09.20.613600

**Authors:** Raza N. Malik, Daniel S. Marigold, Mason Chow, Gevorg Eginyan, Tania Lam

## Abstract

**Background:** Disorders in the recovery of gait strategies in individuals with incomplete spinal cord injury (SCI) suggest difficulties in controlling lower limb intersegmental dynamics, which could relate to proprioceptive impairments. To probe discrete aspects of lower limb interjoint coordination, we present here a novel protocol to assess lower limb motor strategies and evaluate the influence of proprioceptive impairments following SCI.

**Methods:** Twelve able-bodied controls and 16 participants with SCI performed lower limb pointing to three targets that involved combined hip and knee flexion, or hip or knee flexion only while standing, with either full or obstructed visual feedback. We quantified lower limb proprioceptive sense in individuals with SCI using a robotic gait device. We used motion analysis to determine lower limb joint angles and foot trajectory, computed inverse dynamics to quantify joint and intersegmental dynamics, and derived muscle torque as an indicator of the motor strategies produced to control the motion to each target. We used linear mixed-effects models to assess differences between the control and SCI groups on end-point performance and muscle torque, and to assess the relationship of muscle torque with end-point performance and proprioceptive sense.

**Results:** Groups differed in motor strategies, but not end-point performance, when pointing to all three targets. Compared to controls, the SCI group had difficulty controlling knee muscle torque when performing the hip-flexion-only target (p = 0.008) or when flexing the hip and knee simultaneously (p = 0.0004). To complete the knee-flexion-only target, the SCI group had difficulties generating the required hip extensor muscle torque to maintain the thigh in neutral (p = 0.0001). These altered motor strategies in individuals with SCI were associated with proprioceptive impairments and end-point performance.

**Conclusion:** This novel lower limb pointing task can identify disordered motor strategies in individuals with SCI, especially at the knee, and are associated with proprioceptive impairment. Variations of this paradigm can be employed to further understand differences in motor strategies between controls and individuals with SCI, and the impact of proprioceptive deficits.

## Introduction

Locomotor control required to maneuver in our everyday environments relies on precise interjoint coordination to control the simultaneous rotation of the lower limb segments. The control of our legs, which are multi-segmental structures, requires the nervous system to generate precisely timed muscle activation patterns to produce movement and control for passive torques [1–3]. Passive torques, such as gravity and interaction torques, arise from the velocity and acceleration of concurrently moving segments [4]. Biomechanical studies demonstrate that when we step over an obstacle, interaction torques facilitate the required increase in flexion to clear the obstacle [5]. Knee flexion is primarily controlled by active contraction of the knee flexor muscles [5–7], whereas hip and ankle flexion are facilitated via intersegmental dynamics, specifically the passive movement-dependent torques arising from knee linear acceleration and shank angular acceleration [5]. A variety of neurological conditions, including spinal cord injury (SCI), can disrupt this control.

Gait kinematic data of individuals with SCI indicate that they primarily rely on increasing knee flexion to step over obstacles, with little modulation in hip flexion angle, and in some cases even hiking the hip to clear the obstacle [8]. Rehabilitation studies have indicated that individuals with SCI may have general difficulties in modulating interjoint coordination during gait [9–11]. Limitations in the ability to change interjoint coordination following training may be associated with poorer recovery of skilled walking function [12].

Classic studies of individuals with large-fiber neuropathy, resulting in complete proprioceptive loss, have provided important insights into the contribution of proprioceptive sense on interjoint coordination and the control of intersegmental dynamics [13,14]. Sainburg et al. (1995) created a reaching task requiring a series of out-and-back reaching movements to various targets with and without visual feedback of the upper limb. Reaching to different targets requiring varying levels of shoulder-elbow coordination revealed that a smooth reversal during the out-and-back trajectory depends on elbow and shoulder joint movement changing direction synergistically [13]. Moving to targets requiring synchronous shoulder and elbow flexion involved more complex coordination of interaction torques compared to movements to targets requiring a simple, single joint movement, such as isolated elbow flexion. Able-bodied controls moved smoothly and accurately by scaling the amplitude and timing of muscle activity to the magnitude of interaction torques, irrespective of limb visual feedback. In contrast, individuals with large-fiber neuropathy used visual information to partially compensate for their diminished proprioceptive sense and control of intersegmental dynamics to maintain reach accuracy and trajectory [13,15,16]. In the absence of vision, movements in individuals with large-fiber neuropathy became uncoordinated, spatially distorted, and less accurate, especially to targets which involved greater intersegmental dynamics.

Individuals with SCI have varying degrees of impairments in proprioceptive sense, assessed by either joint position sense [17] or movement detection (kinesthetic) sense [18]. Deficits in these aspects of proprioceptive sense can impact the performance of skilled walking [19,20]. When stepping over an obstacle, able-bodied controls (with no proprioceptive deficits) can maintain consistent foot-obstacle clearance height despite varying amounts of peripheral visual feedback from the lower visual field [19,21,22]. However, individuals with SCI increase their foot-obstacle clearance if their lower visual field is blocked when crossing an obstacle, and this change in foot-obstacle clearance height is related to the degree of lower limb joint position sense (but not movement detection sense) [19]. In another study where participants had to perform a foot-target matching task during treadmill walking, we also found that performance error in individuals with SCI associated with the degree of impairment in joint position sense [20]. The associations between proprioceptive impairments and foot-control in individuals with SCI points to impaired control of intersegmental dynamics.

In this study, we aimed to create a novel lower limb pointing task, analogous to the reaching paradigm described above [13,14], to measure intersegmental control and the impact of proprioceptive impairments (both position and movement sense) after SCI. This leg pointing task, instead of a walking or stepping task, allowed us to minimize the effects of potential confounders to intersegmental coordination during locomotion, such as dynamic balance and inter-limb coordination, while enabling us to control for variations in strength and flexibility across individuals. Our pointing task systematically manipulated lower limb interjoint coordination by varying the location of an end-point target in the sagittal plane, as this is the primary plane of body progression during walking. We did not restrict any segment during leg pointing, inevitably resulting in motion of all three segments and joints when pointing to the different targets. Reaching studies where the upper limb is allowed to move unrestricted and against gravity highlight the importance of investigating motor strategies in unrestrained conditions, as strategies differ depending on the effects of gravity and whether limb segments are restricted [23,24]. Here, we used muscle torque, derived from inverse dynamics, to represent motor strategies of the lower limb. When examining the control of intersegmental dynamics, net joint muscle torque can indicate the active elements of the motor strategy (although the effects due to deformation of passive structures cannot be discounted) [25]. To further probe the role of proprioceptive sense, we compared task performance between full and obstructed visual feedback of limb position. Compared to controls, we predicted that individuals with SCI would show disordered end-point (foot) performance and motor strategies when pointing to a target requiring greater interjoint coordination. We expected that limiting visual feedback would exacerbate these effects. We also predicted that greater proprioceptive impairments in individuals with SCI would associate with worse end-point performance and disordered motor strategies.

## Methods

### Participants

We recruited able-bodied controls and individuals with chronic motor-incomplete SCI between the ages of 18-65 years of age to participate in this study. We only included participants in the SCI group if they were at least 1-year post-injury and able to walk at least 5 m, with or without a gait aid. We excluded controls and individuals with SCI participants if they had any other neurological, musculoskeletal, visual, or cognitive deficits affecting mobility. All participants provided written informed consent, and the University of British Columbia Clinical Research Ethics Board approved all procedures.

### Protocol

#### Baseline assessments

We characterized participants by their walking function, muscle strength, and proprioceptive sense. We assessed walking function with the 10-meter walk test, a valid and reliable measure of walking speed in people with SCI [26,27]. We quantified walking speed as the average time (two trials) it took participants to walk 10 meters at their self-selected and maximum speeds [28]. We tested muscle strength for each leg using the lower extremity motor score (LEMS) of the American Spinal Cord Injury Association (ASIA) Impairment Scale (AIS) [29]. Higher scores indicate greater muscle strength.

We quantified proprioceptive sense using joint position sense (JPS) and movement detection sense (MDS) at the hip and knee joints of the right and left legs using previously validated custom-written software of the Lokomat [17,18]. Briefly, participants were suspended in the Lokomat with the non-test limb supported on an elevated surface and the test limb unsupported and lifted off the ground. We used a drape to occlude visual feedback of the limbs. We tested JPS by first moving the test joint (hip or knee) into 25 degrees of flexion or extension and instructed the participant to memorize this position. Following a short delay, the Lokomat moved the test joint to a distractor position. The participant then used a joystick to move the test joint back to the memorized position, and we recorded this angle (further details in *Data Analysis*). We collected six trials at the hip and knee joints for both the right and left legs.

To test MDS, the Lokomat positioned the test joint (hip or knee) in a slightly flexed position. After a variable delay, the Lokomat moved the test joint into flexion or extension at one of three speeds (0.5, 1, or 2 degrees/sec). We instructed the participant to press a button on the joystick as soon as they detected movement in any direction and to verbalize the direction of movement (flexion or extension). We collected 11 trials at the hip and knee joints for both the right and left legs. Specifically, we collected 5 trials with the test joint moving into flexion, 5 trials with the test joint moving into extension, and 1 trial when the test joint did not move (catch trial).

#### Task Setup

Participants stood between parallel bars with a large projection screen (2.21 m wide and 1.60 m high) positioned ∼6 m in front of them (Fig. 1A). We placed rigid bodies, composed of four motion capture markers each, on the thorax and bilaterally on the thighs, shanks, and feet. We also placed individual markers bilaterally on the greater trochanter, lateral femoral condyle, lateral malleolus, and the tip of the big toe. We collected motion capture data at a sampling frequency of 65 Hz (Optotrak, NDI, Waterloo, ON).

**Fig. 1.**
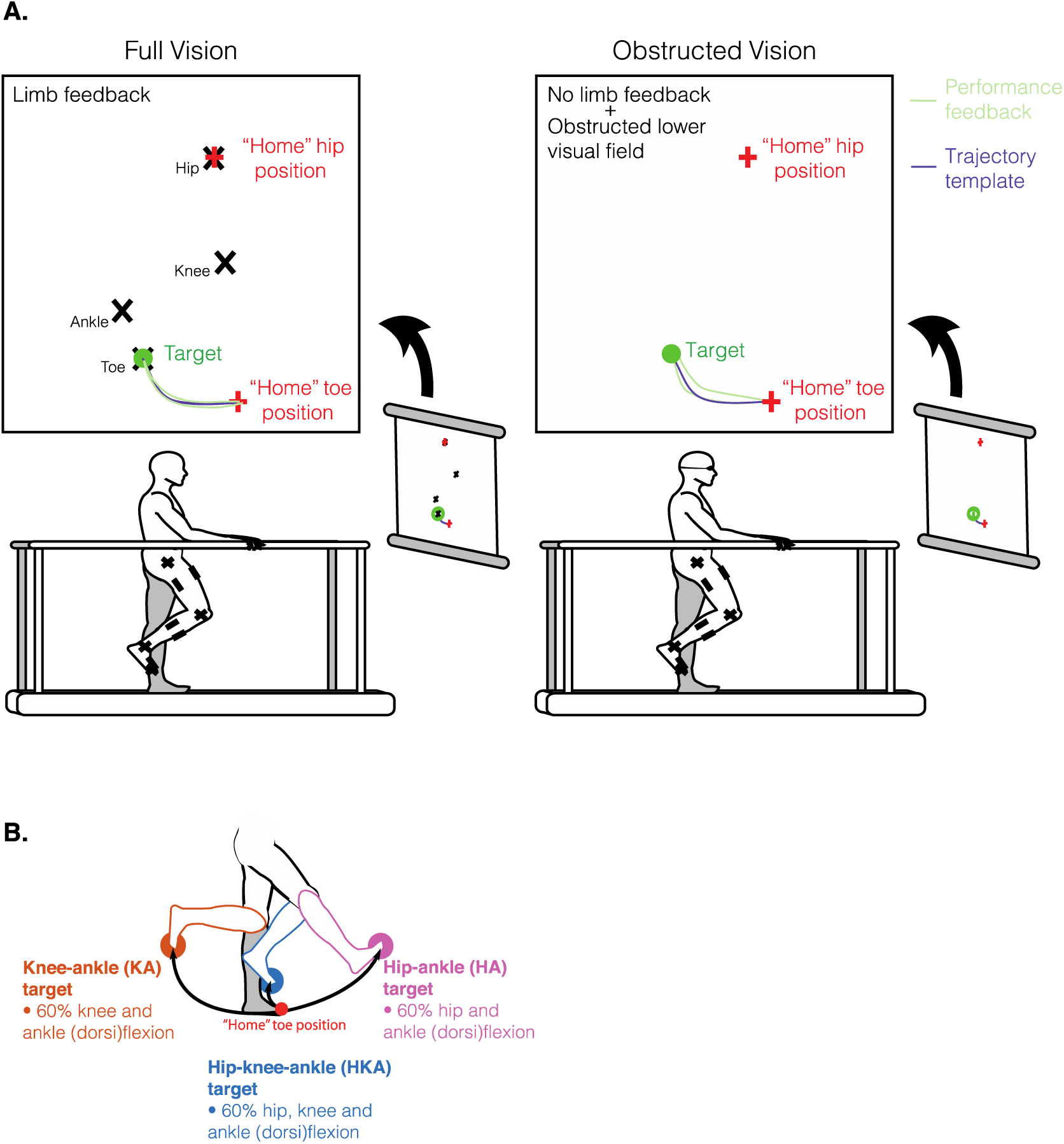
Experimental setup and procedures. A. Participants stood between parallel bars facing a projector screen. We placed motion capture markers on the pointing leg of the lower limb. In the full vision condition, participants could see the position of their hip, knee, and ankle on the screen (black X’s). Before initiating a trial, participants needed to match their hip and toe positions to the “home” hip and toe positions; this ensured participants had a similar starting position across trials. Once they achieved the home position, we verbally cued participants of the upcoming trial. After a short delay, the target and toe trajectory template appeared on the screen. Participants needed to point with their toe to the target while tracing the template in one smooth motion, as accurately as possible. In the obstructed vision condition, participants wore dribble goggles (obstructing the lower visual field), and we did not give joint feedback during pointing. All other parameters were the same as the full vision condition. B. Participants pointed to three different targets in this study, indicated by the different colours. The targets comprised of different combinations of hip, knee, and ankle (dorsi)flexion to 60% of each joint’s maximum range of motion.

We then recorded each participant’s active range of motion at the hip, knee, and ankle of the pointing limb. The pointing limb for the SCI participants was the limb with greater proprioceptive impairments. In controls, the pointing limb was the non-dominant limb, as determined by the Waterloo Footedness Questionnaire [30]. We asked participants to flex their hip only, knee only, or dorsiflex their ankle only from a standing position while we recorded the maximum angle achieved at each joint during these motions. We recorded two trials for each joint and averaged them to determine the active range of motion at the hip, knee, and ankle joints. In the control group, the mean (SD) maximum range of motion at the hip, knee, and ankle was 64.7° (18.5), 103.1° (12.1), and 17.1° (6.3), respectively. In the SCI group, the mean (SD) maximum range of motion at the hip, knee, and ankle was 47.5° (10.0), 86.6° (17.8), and 10.5° (8.2), respectively.

Using Unity (Unity Technologies, San Francisco, USA), we streamed the real-time signals from the Optotrak markers over the greater trochanter, lateral femoral condyle, lateral malleolus, and big toe (representing the hip, knee, ankle, and toe) of the pointing limb, and presented the signals on the projection screen. We displayed the hip, knee, ankle, and toe sagittal-plane positions as X’s forming a virtual limb on the projection screen (Fig. 1A). The virtual limb only responded to participant movement in the sagittal plane where anterior-posterior limb movements resulted in the virtual limb moving to the right and left, respectively, on the screen. Cranial-caudal limb movements resulted in the virtual limb moving up and down, respectively. Once the participant familiarized themselves with the virtual limb, we asked them to stand in a comfortable upright posture. We captured the position of the hip and toe and displayed them as red ‘+’ signs; this was the participant’s “home position” (Fig. 1A) and was used to ensure that participants began each trial from the same position.

We created target positions and toe trajectory templates in Unity (see below). The target positions were determined by different combinations of hip, knee, and ankle joint flexion scaled to 60% of each joint’s active range of motion. In this way, we could normalize the target positions across participants, accounting for inter-individual differences in strength and flexibility. We also generated the optimal toe trajectory path to each target by modelling the participant’s limb and rotating the appropriate joint(s) (for that target) to 60% of their active range of motion (*trajectory template*, Fig. 1A). For example, the toe trajectory template and target location when pointing to the hip-ankle (HA) target would result from moving only the hip and ankle through 60% of their full joint range of motion from the home position, with no movement at the knee.

The different combinations of joint flexion resulted in a total of seven targets that comprised of both single jointed (e.g., hip or knee target) and multi-jointed targets (e.g., hip-knee or hip-knee-ankle target). In both the control and SCI groups, there were no statistical differences (linear mixed-effects model, p > 0.05) between pointing to targets that included the ankle (hip-ankle target, knee-ankle target, hip-knee-ankle target) compared to targets that did not account for ankle motion (hip-only target, knee-only target, hip-knee target) for all outcome measures, except for vertical end-point error. Pointing to the non-ankle targets increased vertical end-point error by 1.16 cm in the controls (p < 0.001) and by 1.05 cm in the SCI group (p < 0.001) compared to pointing to targets that included the ankle. Given that most outcome measures were not statistically different between targets that included the ankle and those that did not, and for simplicity of the analysis, we combined the ankle and non-ankle data from corresponding targets. Therefore, we combined hip-only with hip-ankle, knee-only with knee-ankle, and hip-knee with hip-knee-ankle targets to simplify the analysis to three targets: hip-knee-ankle (HKA), knee-ankle (KA), and hip-ankle (HA) targets (Fig 1B).

#### Pointing task

We instructed participants to point (using their toe) as accurately as possible to one of the targets described above, projected onto the screen in front of them (Fig. 1A). Once participants were in their home position, a verbal cue indicated the start of the trial. The target and toe trajectory template appeared on the projection screen after a short delay (∼3-5 s) (Fig. 1A). We instructed participants to move to the target, then return to the home position in one smooth motion, following the toe trajectory template (*Trajectory template,* Fig. 1A). After each trial, we provided visual feedback of their toe trajectory on the screen (*Performance feedback,* Fig. 1A). We presented targets during full and obstructed vision conditions. During full vision trials, the participant’s visual field was unobstructed, and the virtual limb was visible on the projection screen throughout the movement (*Full Vision*, Fig. 1A, left panel). During obstructed vision trials, we obstructed the participant’s lower visual field using dribble goggles and removed feedback of the virtual limb on the projection screen during the movement (*Obstructed Vision*, Fig. 1A, right panel). We gave visual feedback of toe trajectory on the projection screen after every trial, irrespective of visual condition.

Prior to starting the task, we familiarized participants with each target. During familiarization, we informed participants of the target to be presented and the joint motion required; participants received 3-5 blocked trials with toe trajectory feedback at each target. For example, when we presented the HA target, we informed the participant that this was the “hip-ankle target”, and that pointing to it required that they flex their hip and ankle only, with no movement at the knee. We did block familiarization for all targets ensuring the participant was comfortable with the virtual limb and understood the task and how to point to all targets. After familiarization with each target, we presented the targets in random order in two blocks of 42 for a total of 84 trials. Six trials at each target were presented randomly per block. We administered the first block with full vision and the second block under obstructed visual conditions.

### Data Analysis

We characterized proprioceptive sense by *JPS* and *MDS*. We calculated JPS, in degrees, as the absolute mean difference between the target and actual position for six trials at each joint. Higher scores indicate poorer JPS [17]. We calculated MDS, in arbitrary units (a.u.), by taking the sum of the joint excursion before the button was pressed divided by 10 (the maximum possible joint angle) and the participant’s response to the direction of movement (0 = correct response, 1 = incorrect response). Higher scores (up to a maximum of 2) indicate poorer MDS, and lower scores (to a minimum of 0) indicate better MDS [18].

We derived toe trajectory and joint angles in Visual 3D (C-Motion, Inc., Rockville, MD) based on each participant’s lower limb model. We exported and analyzed these kinematic variables in MATLAB (MathWorks, Natick, MA). We low-pass filtered the kinematic data with a 4^th^-order dual-pass Butterworth filter at 6 Hz. For each trial, we focused our analysis to the reversal phase, which requires precision control of the different joints to control foot trajectory. We defined the *reversal phase* as the movement between the peaks in the magnitude of toe velocity over the entire movement [13,31] (blue shaded region, Fig 2A). The *reversal point* occurs when toe velocity is at its lowest during the reversal phase, indicating the time at which the toe reverses direction (Fig 2A, black dot). The period from the first toe velocity peak (Fig 2A, red dot) to the reversal point indicates movement towards the target (Fig 2A, thick red line). The period between the reversal point to the second toe velocity peak (Fig 2A, blue dot) indicates movement away from the target, back towards the start position (Fig 2A, thick blue line). Thus, the reversal phase corresponds to toe deceleration when moving to the target and toe acceleration when moving back towards the start position [13,31]. We used toe position at the reversal point to calculate end-point error (Fig 2B, black dot) and peak segment angles (Fig 2C). We used toe trajectory over the entire reversal phase to calculate foot-path area (Fig 2B, checkered area).

**Fig. 2:**
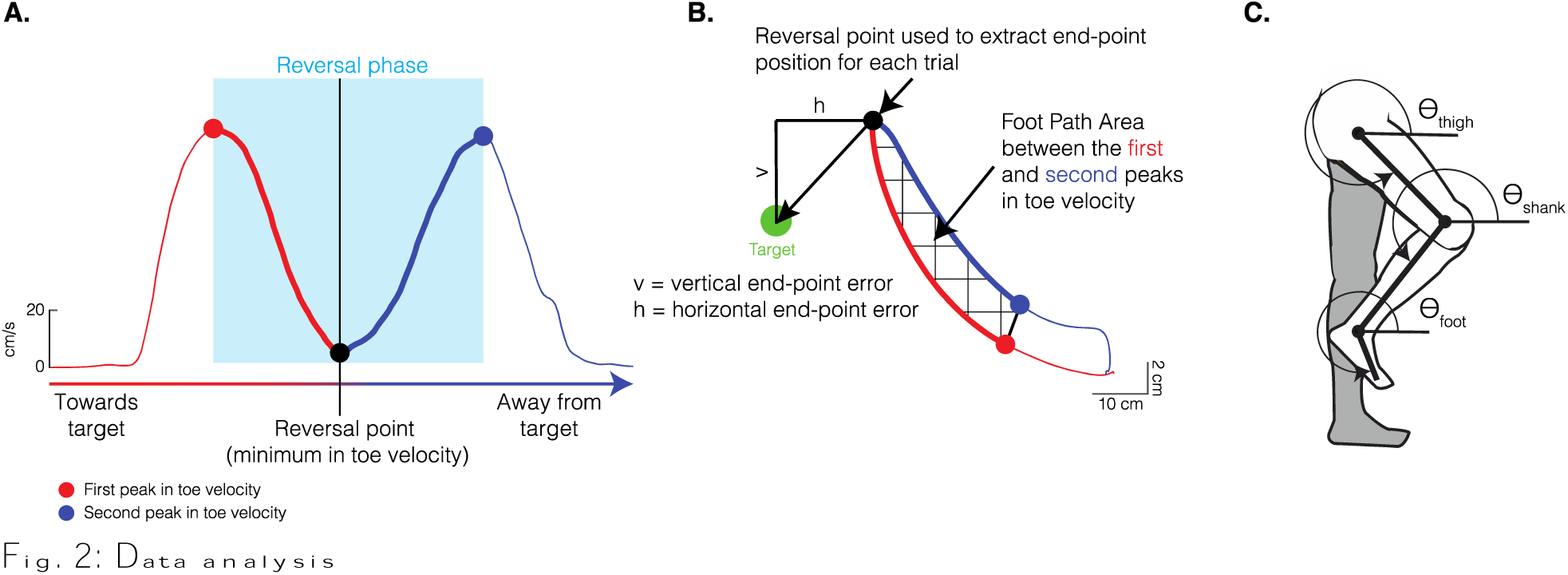
Data analysis. A. Toe velocity profile from a single trial. We used the peaks (red and blue dots) and valley (black dot) of the toe velocity signal to characterize the reversal phase of the movement. This phase consisted of an initial deceleration towards the target (thick red line) followed by acceleration away from the target back toward the start position (thick blue line). B. Toe trajectory profile of data shown in A. We used toe trajectory to calculate measures of end-point accuracy. We used toe position at the reversal point (black dot) as the end-point position for each trial. We then calculated the horizontal (h) and vertical (v) end-point errors as the vertical and horizontal distance of the end-point relative to the target. We calculated foot-path area by determining the area encompassed by the toe during the reversal phase (checkered area). The red and blue dots indicate the first and second peaks in toe velocity, respectively, and define the reversal phase (see A). C. We used a three-segment (thigh, shank, foot) lower limb model that was interconnected via frictionless joints (hip, knee, ankle). We calculated segment angles relative to the right horizontal. Segment angles and their derivatives (velocity and acceleration), alongside anthropometric measurements, were used in the equations of motion to calculate joint torques [32]. We quantified joint torques over the first half of the reversal phase (thick red line in A).

Joint torques over the first half of the reversal phase (Fig. 2A, thick red line) were used to quantify motor strategies. To control for possible effects of movement speed on intersegmental dynamics (see Statistical Analysis), we calculated movement time as the time (in seconds) it took to complete the first half of the reversal phase. Overall, across targets and vision conditions, the control group’s mean movement time was 0.64 (SD 0.23) seconds and that of the SCI group was 0.58 (SD 0.22) seconds.

### End-point error and foot-path area

We investigated end-point (toe) control by examining *end-point error* along the vertical and horizontal axes and foot-path area. We calculated end-point error for each trial as the vertical and horizontal distances between toe position at the reversal point and the vertical and horizontal position of the target, respectively (Fig. 2B). Negative horizontal end-point error values indicate that the participant’s toe position at the reversal point was posterior to the target, and positive error values indicate that toe position was anterior to the target. Thus, ‘overshooting’ the KA and HA targets along the horizontal axis corresponded to posterior and anterior end-point errors, respectively, and vice versa for ‘undershooting’. For vertical end-point error, negative values indicate undershooting the target (for all targets) and positive values indicate overshooting in the vertical axis.

We defined *foot-path area* as the area within the trajectory of the foot during the reversal phase (Fig. 2B, checkered area). We divided the reversal phase into two halves to calculate foot-path area: 1) the start of the reversal phase (Fig. 2B, red dot) to the reversal point (Fig. 2B, black dot), and 2) the reversal point to the end of the reversal phase (Fig. 2B, blue dot). We calculated the definitive integral under each half using the trapezoid rule to determine the area under each half. We defined foot-path area as the area between the two halves by taking the difference between the definitive integrals of each half. Larger values indicate a larger foot-path area.

### Motor strategies: Interaction and muscle torques

We used inverse dynamics and equations of motion to investigate dynamic interactions between segments [32]. We used a lower extremity model of three segments (thigh, shank, foot) connected by three frictionless joints (Fig 2C). We derived segment angles with respect to the right horizontal (Fig 2C), which were then used to derive segment angular velocities and accelerations. We extracted bilateral segment masses, lengths, center of masses, and moments of inertia from Visual 3D. We used the kinematic and anthropometric measurements in the equations of motion, as described in Hoy and Zernicke (1986), to calculate net, interaction, gravitational, and muscle torques. Following the model of Hoy and Zernicke (1986), we further parsed the interaction torques into those arising from the angular velocity and acceleration of the thigh, shank, and foot segments; torque components due to hip linear acceleration was set to zero. We derived muscle torque by subtracting the net torque from the sum of the gravitational and interaction torques [32]. The generalized muscle torque derived here includes forces generated from active muscle contractions, as well as passive deformation of muscles, tendons and ligaments [32].

We focus on *joint muscle torque* to represent motor strategies used by the participants to control intersegmental dynamics [13,25,31,33]. To quantify muscle torque, we calculated its mean magnitude between the first peak in toe velocity and the reversal point (darker shaded blue, Fig 2A). Quantifying joint muscle torque during this period allowed us to examine participants’ control strategies to point to the target (and not those for returning to the home position). We also calculated *muscle power* (the product of the muscle torque and relative angular velocity at each joint), to describe the mechanical work performed at each joint [6,34,35].

### Statistical Analysis

We used RStudio Version 4.02 for all statistical analyses. We used descriptive statistics, including mean, median, confidence intervals and standard deviation, frequency, and percentages, as appropriate, to summarize participant characteristics, end-point performance (horizontal and vertical end-point error, foot-path area), segment angles, and motor strategies (torque profiles). The outcome variables entered into the statistical models below included the continuous variables of muscle torque, vertical and horizontal end-point error, and foot-path area. Our primary outcome variable was muscle torque, which indicated the motor strategies used by the participants. Secondary outcome variables were the measures of end-point performance (vertical and horizontal end-point error, and foot-path area), which indicate the performance of the task arising from the motor control strategy.

We fitted linear mixed-effects models (LMEs) to analyze the relationship between outcome and predictor variables while accounting for repeated observations per participant. We used a random intercept model for all LMEs that included categorical and continuous predictor variables. We modeled the LMEs in R [36] using the lme4 package[37]. We created and visualized tables of estimates, confidence intervals, p values, and random effects using the sjPlot package [38]. We assessed model fit using information criteria (Akaike Information Criteria (AIC)), where smaller values represent a better fitting model [39]. We used the Wald Test and AIC to identify the best fitting model. All models were estimated using Maximum Likelihood.

To estimate differences in motor strategies and end-point performance between groups, we generated LMEs for each outcome variable that included the fixed effects of target (HKA, KA, HA), vision (full, obstructed), and group (control, SCI), and the random effect of participant. The referents for target, vision, and group were the HKA target, full vision, and control group, respectively. We controlled for maximum joint range of motion and movement time by including these continuous variables in the model. We chose our final model by first including the interaction terms assessing the two-way interaction between target and group, and between vision and group. A three-way interaction was only assessed if both two-way interactions were statistically significant. If no two-way interactions were observed, the predictors were entered into the model without interacting with other predictors.

We further evaluated meaningful interactions with pairwise comparisons using the emmeans package [40]. When group differences or interactions in motor control strategies were found, we first sought to further explore the impact of altered motor control strategies in the SCI group on task performance. We did this by evaluating the crude associations between motor strategy (muscle torque) and the three end-point performance variables (vertical and horizontal end-point error, and foot-path area) in the SCI group using LMEs, where an end-point performance variable was entered as the outcome variable and muscle torque as the predictor variable.

To estimate the impact of lower limb proprioceptive deficits on end-point performance and motor strategies in the SCI group, we focused only on the targets and those outcome variables where we found group differences or interactions in our first LME analysis using crude associations between those outcome variables and proprioceptive sense.

We conducted model diagnostics for all LMEs using the DHARMa [41] package on each outcome’s final model. We inspected the QQ (quantile-quantile) plot of residuals and a plot of residuals against predicted values to test the assumptions of normality and homogeneity of variance. Foot-path area was the only variable that we log-transformed to meet the assumption of normality. We back-transformed this data for visual presentation.

We present effect sizes here to facilitate future *a priori* sample size calculations for studies based on the protocol presented here. We used Cohen’s *f*^2^ to determine the effect size of the variance explained by each model (marginal R^2^) [42]. We used Cohen’s *d* to determine the effect size of estimated mean group differences for the pairwise comparisons.

## Results

All participants who enrolled in the study were able to complete the full study protocol. The data presented in this study are from 12 controls (5 males and 7 females; mean age = 41 years (range 20-65); mean height = 170 cm (range 151-186); mean weight = 71 kg (range 45-98)) and 16 individuals with motor-incomplete SCI (10 males and 6 females; mean age = 54 years (range 28-66); mean height = 169 cm (range 149-185); mean weight = 75 kg (range 50-109)).

In controls, mean MDS and JPS scores were 0.2 a.u. (range 0.1-0.4) and 6.63° (range 2.33-10.27), respectively. In the SCI group, the mean MDS and JPS were 1.03 a.u. (range 0.13 – 2.43) and 12.29° (range 4.07-35.8), respectively. Both MDS and JPS were significantly worse in the SCI group compared to controls (*t-test*, p < 0.05). Detailed participant characteristics of the SCI group are shown in Table 1.

**Table 1.**
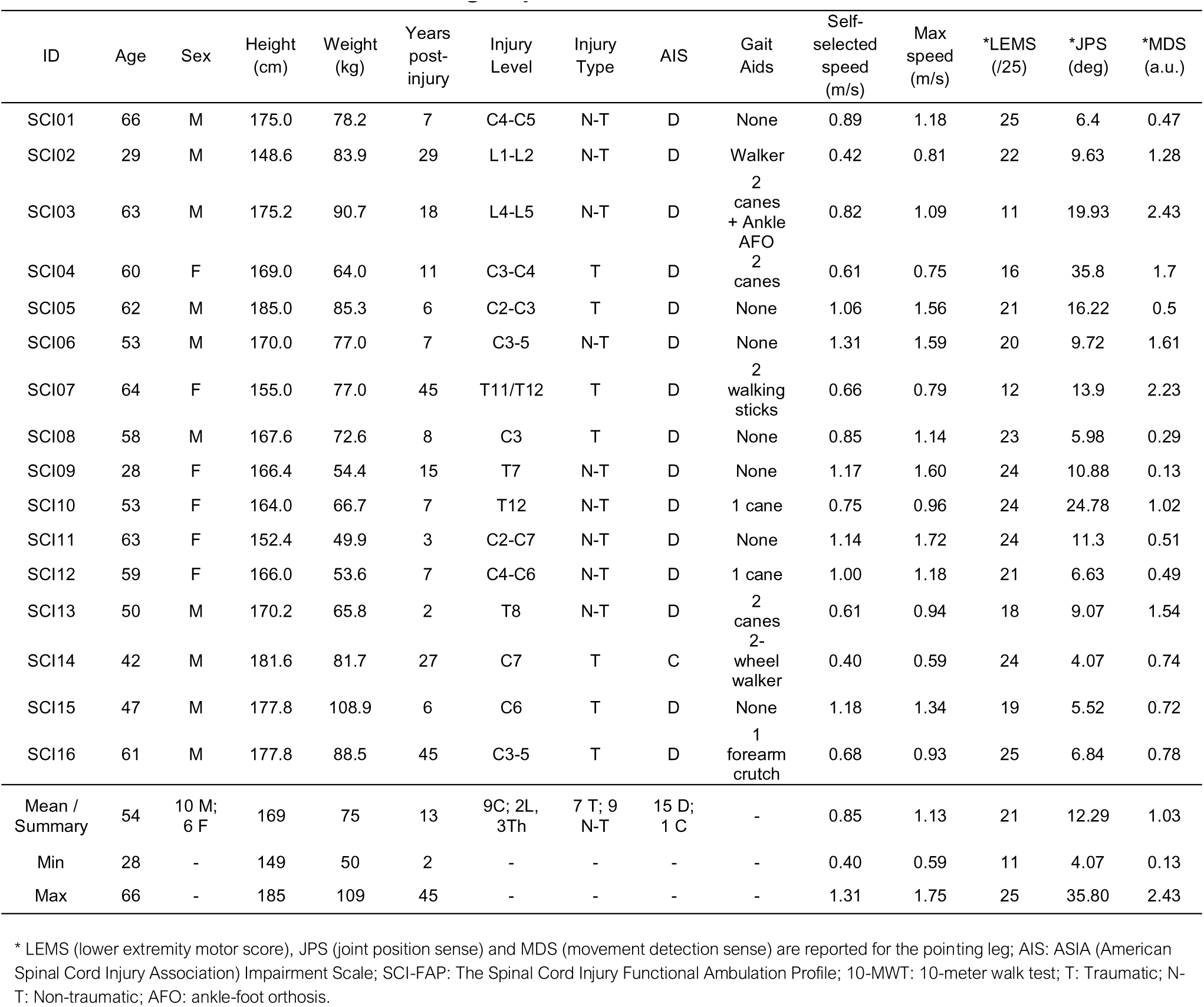
Characteristics of the SCI group.

### End-point performance

Figure 3 shows toe trajectories from a representative control and two SCI participants (with contrasting proprioceptive sense) pointing to the different targets in both visual conditions. Comparing the trajectory profiles of the control participant and SCI01 (who both had comparable proprioceptive sense) with those of SCI04, we see that SCI04 demonstrated greater difficulty in controlling toe trajectory, possibly related to their poorer proprioceptive sense.

**Fig. 3.**
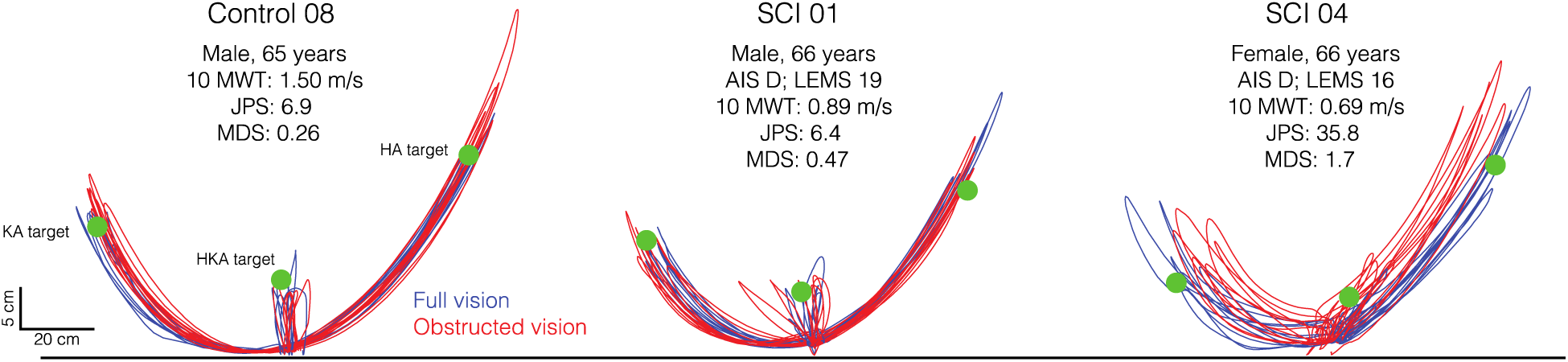
Toe trajectories. Toe trajectories from a control participant and two individuals with spinal cord injury (SCI) when pointing to the different targets under full (blue) and obstructed vision (red). Relevant characteristics (e.g., proprioceptive sense) are reported for each participant. 10-MWT: 10-meter walk test; JPS: joint position sense; MDS: movement detection sense; AIS: ASIA (American Spinal Cord Injury Association) Impairment Scale; LEMS: Lower Extremity Motor Score

In the full vision condition, controls could perform the task with minimal horizontal end-point error to all three targets (Fig. 4, black symbols). There was minimal vertical end-point error at the HKA target but larger errors at the HA and KA targets. We also found the largest foot-path area at the HKA target and smaller area at the KA and HA targets. Generally, end-point errors were larger during obstructed vision, while foot-path area showed little difference between vision conditions.

**Fig. 4.**
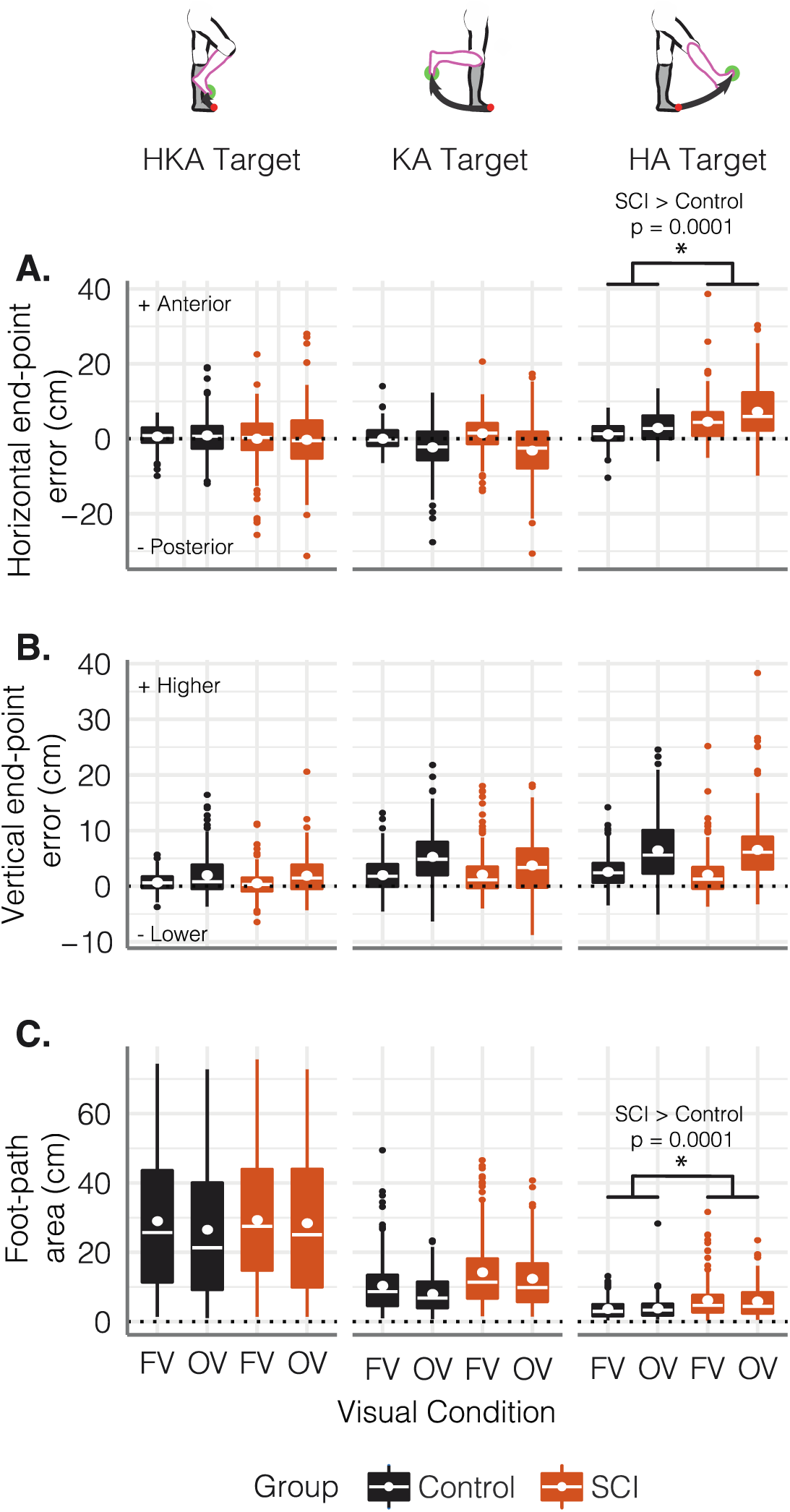
End-point performance when pointing to the different targets. Boxplots depicting A. horizontal end-point error, B. vertical end-point error, and C. foot-path area in the control (black) and SCI (orange) groups in both visual conditions (FV: full vision; OV: obstructed vision) across all three targets. Asterisks represent significant post-hoc group differences between controls and the SCI group averaged across visual conditions. Post-hoc comparisons were conducted after detecting a significant group x target interaction in the linear mixed-effect models (see text for full statistical details).

For end-point performance, the LME results suggest that target modifies group effects for horizontal end-point error (Fig 4A; *LME:* group × target interaction, p < 0.001, marginal R^2^ = 0.141, f^2^ = 0.164) and foot-path area (Fig 4C; *LME:* group × target interaction, p < 0.001, marginal R^2^ = 0.434, f^2^ = 0.767). There were no group main effects or interactions for vertical end-point error. We only detected post-hoc group differences during the performance of the HA target (Fig 4, panels with asterisks). Compared to controls, our sample of individuals with SCI had larger anterior errors (over-shooting) (estimated difference: 4.125 [95%CI: 2.25, 6], t(45.2) = 4.431, p = 0.0001, *d* = 0.71) and larger foot-path areas when performing the HA target (estimated difference: 0.665 [95%CI: 0.521, 0.848], t(46.7) = 3.377, p = 0.0015, *d* = 0.52), irrespective of visual condition.

### Joint torques

Pointing to the HKA, KA, and HA targets may require different motor strategies, as shown by the adjustments observed in segment angles, joint torques, and joint power across the three targets. Figure 5 highlights these motor strategies, as exemplified by the control group (ensemble mean; all movements performed with full vision). Moving to the HKA target (Fig. 5A) involved the production of hip and knee flexor muscle torque and ankle dorsiflexor muscle torque. At the reversal point, the thigh segment reached a rotation of 299.1° (SD 9.8), the shank rotated to 255.6° (SD 13.8), and the foot to 341.2° (SD 15.6). Positive power was generated at all three joints, but the work was primarily done by the hip and knee joints. Interaction torques experienced at the hip and ankle were small but contributed to (dorsi)flexion, thereby assisting muscle torque in counteracting gravity. The interaction torque experienced at the knee was also small but contributed to extension, as did the gravitational torque; both were counteracted by the muscle torque. The interaction torques produced at each joint when pointing to the HKA target primarily arose from hip angular acceleration.

**Fig. 5.**
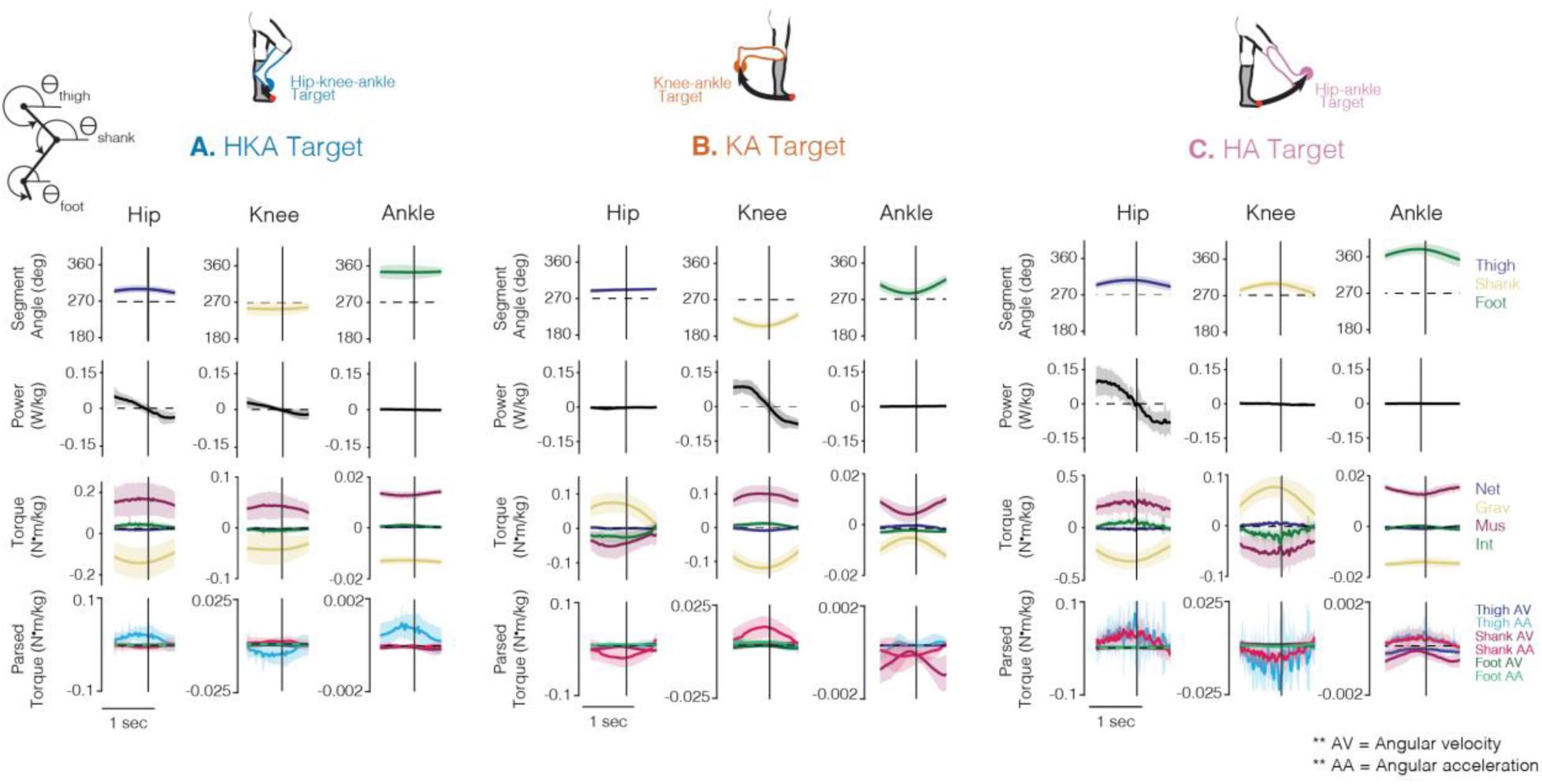
Kinematics, kinetics, and energetics of the lower limb pointing task. Segment angles, joint power, joint torque, and parsed joint interaction torques for the controls (ensemble average, n = 12) when pointing to the A. HKA, B. KA, and C. HA targets in the full vision condition. Shaded areas around each trace represent the 95% confidence intervals. Data are presented for the duration of the reversal phase and vertical bars represent the reversal point (see Fig. 2A). For the power traces, positive values represent generative and negative values represent absorptive power. Flexor/dorsiflexor torques are positive. The inset stick figure in the top left corner indicates the orientation of segment angles (see also Fig. 2C).

When moving to the KA target (Fig. 5B), the thigh remained near vertical (maintained at 278.1° (SD 4.6) at the reversal point), while the shank rotated clockwise from vertical to 236.9° (SD 9.2), and the foot rotated to 321.2° (SD 9.03). The movement primarily involved knee flexor muscle torque and positive power to rotate the shank segment. At the hip, the extensor muscle torque generated would maintain the thigh segment in a neutral position. At the ankle, dorsiflexor muscle torque was reduced as gravity helped to maintain the foot segment vertical, keeping the ankle neutral. Power was primarily generated at the knee, and accordingly, interaction torques at all three joints primarily arose from shank angular acceleration.

When moving to the HA target (Fig. 5C), the thigh segment rotated to 308.8° (SD 11.40) at the reversal point, the shank to 301.3° (SD 10.4), and the foot to 381.9° (SD 13.5). The movement generated hip flexor and ankle dorsiflexor muscle torque, while muscle torque at the knee was extensor to maintain a straight knee during the movement. At this target, power is generated primarily at the hip and close to zero at the knee and ankle. Interaction torques at the hip, knee, and ankle, primarily arising from thigh and shank angular acceleration, assisted muscle torque in counteracting gravity.

### Motor strategies

Our data indicated group differences in motor strategies at all three targets (Figs. 6A-C, *panels with asterisks*). The hip muscle torque showed an overall group effect, which was modified by target (*LME:* group × target interaction, p < 0.001, marginal R^2^ = 0.766, *f^2^* = 3.27). Post-hoc comparisons indicated that individuals with SCI tended to produce less hip extensor torque compared to controls when pointing to the KA target irrespective of visual condition (Fig. 6A *middle panel*, estimated difference: 0.074 [95% CI: 0.034, 0.109], t(33.9) = 4.307, p = 0.0001, *d* = 1.51). This reduced hip extensor muscle torque in the SCI group corresponded to greater horizontal end-point error anteriorly (Fig. 6D *left panel* ; R^2^ = 0.484, p < 0.001, *f^2^*= 0.938) and larger foot-path areas (Fig. 6D *right panel* ; R^2^ = 0.070, p < 0.001, *f^2^* = 0.075), but not vertical end-point error (Fig. 6D *middle panel* ).

**Fig. 6.**
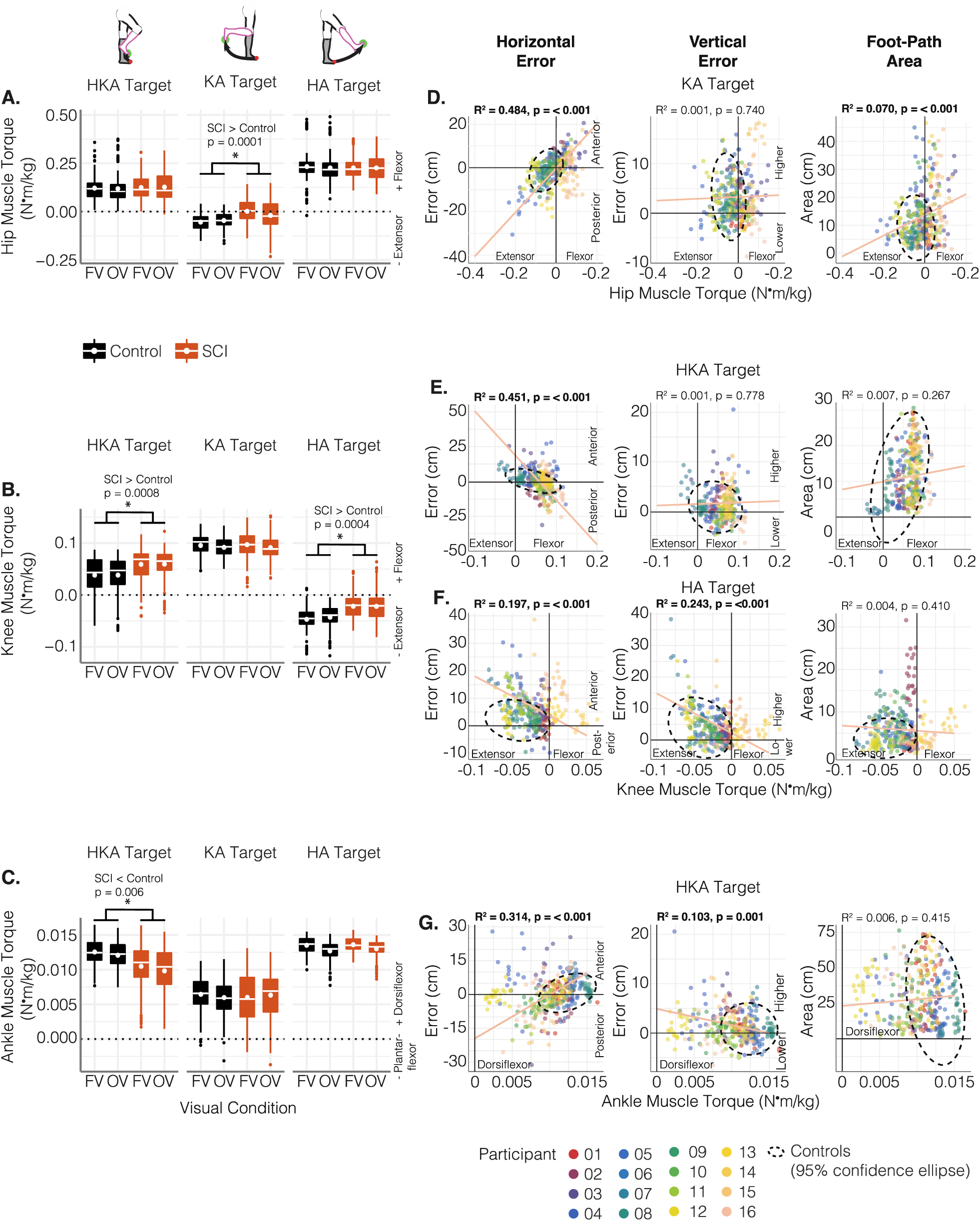
Motor strategies when pointing to the different targets and their associations with end-point performance. Boxplots depicting A. hip muscle torque, B. knee muscle torque, and C. ankle muscle torque in the control (black) and SCI (orange) groups in both visual conditions (FV: full vision; OV: obstructed vision) across all three targets. Asterisks represent significant post-hoc group differences between controls and the SCI group averaged across visual conditions. Post-hoc comparisons were conducted after detecting a significant group x target interaction in the linear mixed-effect model (see text for full statistical details). Scatterplots of D. hip muscle torque when pointing to the KA target, E. knee muscle torque when pointing to the HKA target, F. knee muscle torque when pointing to the HA target, and G. ankle muscle torque when pointing to the HKA target, in relation to horizontal end-point error (left panels), vertical end-point error (middle panels), and foot-path area (right panels) in individuals with SCI. The regression line and R^2^ values were estimated using linear mixed-effect models fitted to assess the crude relationship between the two variables of interest. The black ellipse represents the area encompassing 95% of the control data.

Knee muscle torque also showed an overall group effect that was modified by target (*LME:* group × target interaction, p < 0.001, marginal R^2^ = 0.808, *f^2^* = 4.2). Post-hoc comparisons indicated that individuals with SCI tended to produce more knee flexor muscle torque than controls when pointing to the HKA target (Fig. 6B *left panel*, estimated difference: 0.029 [95% CI: 0.013, 0.044], t(33.7) = 3.701, p = 0.0008, *d* = 1.401), and less knee extensor muscle torque when pointing to the HA target (Fig. 6B *right panel*, estimated difference: 0.030 [95% CI: 0.014, 0.045], t(33.6) = 3.890, p = 0.0004, *d* = 1.472), irrespective of visual condition. This greater knee flexor muscle torque in the SCI group when moving to the HKA target associated with greater horizontal end-point errors posteriorly (undershooting) (Fig. 6E *left panel*; R^2^ = 0.451, p < 0.001*, f^2^* = 0.821) but not vertical end-point error (Fig. 6E *middle panel*) or foot-path area (Fig. 6E *right panel*). The reduced knee extensor muscle torque when moving to the HA target corresponded to smaller horizontal errors anteriorly (Fig. 6F *left panel*; R^2^ = 0.197, p < 0.001, *f^2^* = 0.245) and lower vertical end-point error (undershooting) (Fig. 6F *middle panel*; R^2^ = 0.243, p < 0.001, *f^2^* = 0.321), but did not associate with foot-path area (Fig. 6F *right panel*).

The ankle muscle torque produced when moving to the different targets also showed an overall group effect that was modified by target (*LME:* group × target interaction, p < 0.001, marginal R^2^ = 0.598, *f^2^* = 1.488). Post-hoc comparisons indicated that individuals with SCI tended to produce less dorsiflexor muscle torque at the ankle compared to controls when moving to the HKA target, irrespective of visual condition (Fig. 6C *left panel*, estimated difference: 2.17 x 10^-3^ [95 % CI: 6.53 x 10^-4^, 3.69 x 10^-3^], t(33.3) = 2.909, p = 0.006, *d* = 1.170). This reduced ankle dorsiflexor muscle torque associated with greater horizontal end-point errors posteriorly (Fig. 6G *left panel*; R^2^ = 0.314, p < 0.001, *f^2^*= 0.458) and higher vertical end-point errors (Fig. 6G *middle panel*; R^2^ = 0.103, p = 0.001, *f^2^* = 0.115), but did not associate with foot-path area (Fig. 6G *right panel*).

### Associations between proprioceptive sense and motor strategies

In the individuals with SCI, we investigated the relationship between proprioceptive sense and only those motor strategy variables where the analysis above indicated differences between groups, that is, hip muscle torque when pointing to the KA target (Fig. 6A), knee muscle torque when pointing to the HKA and HA target (Fig. 6B), and ankle muscle torque when pointing to the HKA target (Fig. 6C). However, although our findings above indicated less knee extensor muscle torque in the SCI group compared to controls when pointing to the HA target (Fig. 6B) and less ankle dorsiflexor muscle torque when pointing to the HKA target (Fig. 6C), we did not detect any relationships between proprioceptive sense and these variables. Also, we only detected relationships with MDS, but not JPS. Thus, we only present the relationship between MDS and hip extensor muscle torque when pointing to the KA target and knee flexor muscle torque when pointing to the HKA target.

Representative traces of hip torque while pointing to the KA target in an individual with relatively good (SCI01) and one with relatively poor (SCI04) proprioceptive sense are shown in Fig. 7A. As illustrated in Fig. 6B, normally when moving to the KA target, muscle and interaction torques at the hip are extensor, and work together to oppose gravity in the first half of the reversal phase to keep the thigh segment in a neutral position. In the examples shown in Fig. 7A, there is an apparent disorder in this motor strategy, with both muscle and gravitational torque in the flexor direction, counteracting an extensor interaction torque. Further, among the SCI group, when moving to the KA target, our data suggest that worse MDS was associated with the production of less extensor muscle torque (more flexor muscle torque) at the hip (Fig. 7B; R^2^ = 0.168, p = 0.022, *f^2^*= 0.202).

**Fig.7.**
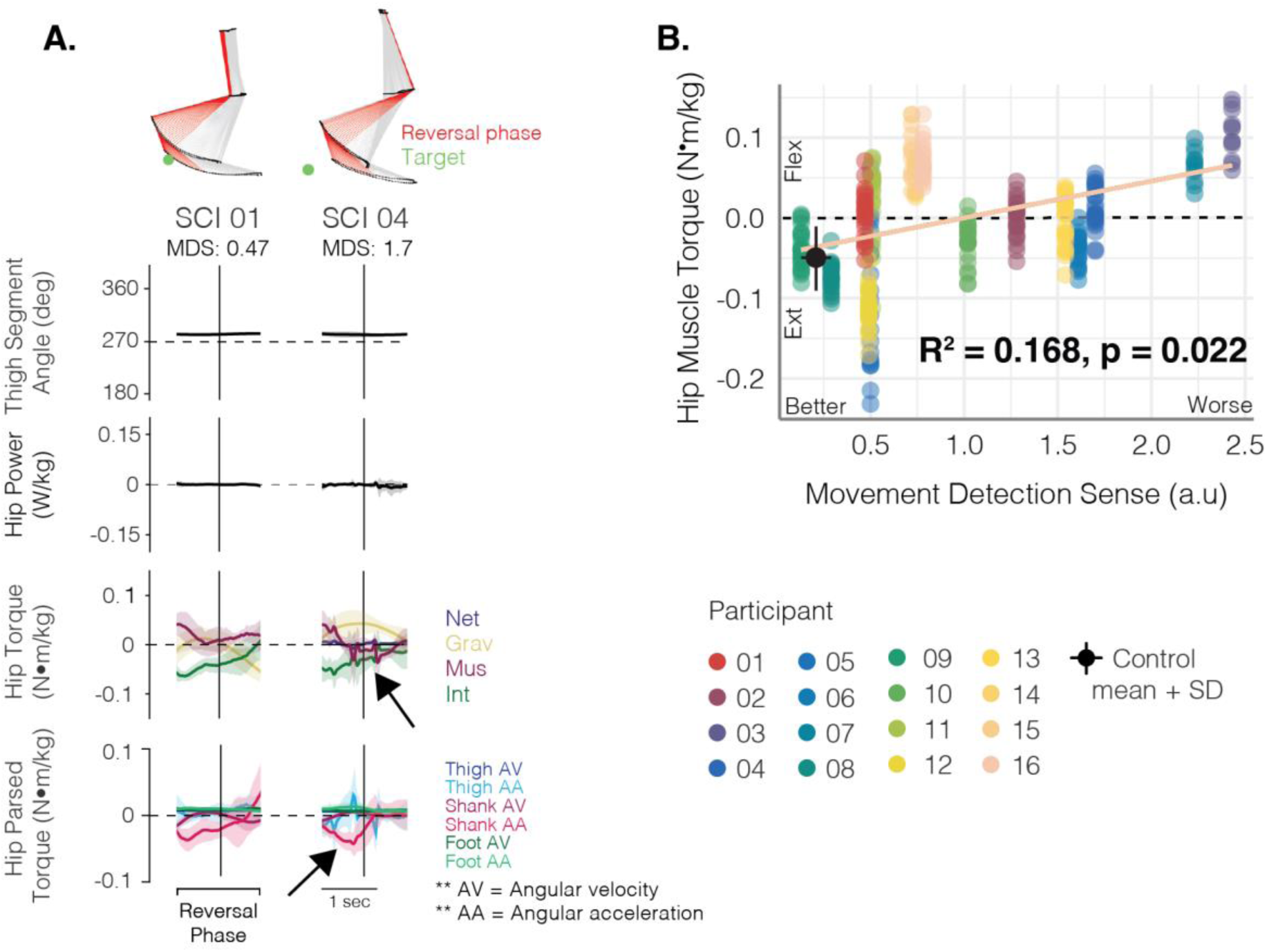
Associations between hip muscle torque and proprioceptive sense at the KA target. A. Thigh segment angles, hip power, and hip joint torque signals for two individuals with SCI when pointing under full vision. SCI04 had worse proprioceptive sense than SCI01. For the power traces, positive values represent generative and negative values represent absorptive power. Flexor torques are positive. Arrows indicate regions of interest highlighting the effects of greater proprioceptive deficits on controlling lower limb dynamics B. Scatterplot of the hip muscle torque produced by each individual with SCI when pointing to the KA target in relation their movement detection sense score, irrespective of visual condition. The regression line and R^2^ value was estimated using a linear mixed-effect model fitted to assess the crude relationship between these two variables. The black dot and error bars represent the mean and standard deviation (SD) of the control data.

Representative knee torque trajectories from SCI01 and SCI04 are shown in Fig. 8A. As illustrated in Fig. 6A, normally when moving to the HKA target, knee torques in the first half of the reversal phase are dominated by a flexor muscle torque that generates power to counteract gravitational torque, with relatively minor contribution from knee interaction torques. In the examples shown in Fig. 8A, knee torque trajectories in SCI01 follow a similar pattern as that observed in controls, while those in SCI04 are greatly reduced, and the accompanying contrast in the amount of knee flexor power is apparent. Accordingly, rotation of the shank during this task is evident in SCI01, while SCI04 appears to hold their shank segment still. Among the SCI group, we noted that when moving to this target, our data suggest that worse MDS was associated with reduced knee flexor muscle torque (Fig. 8B; R^2^ = 0.252, p = 0.006, *f^2^* = 0.337).

**Fig. 8.**
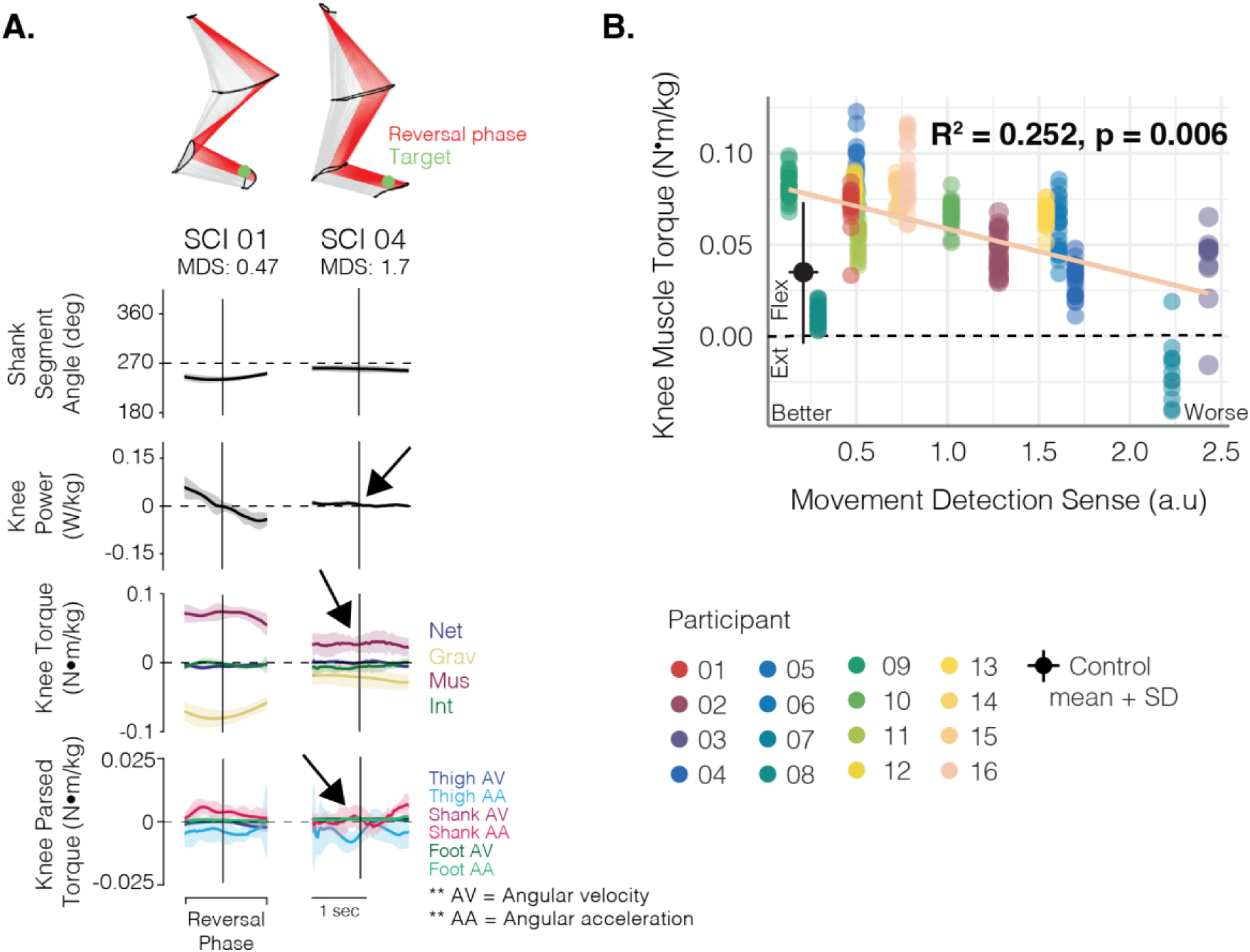
Associations between knee muscle torque with proprioceptive sense at the HKA target. A. Shank segment angles, knee power, and knee joint torque signals for two individuals with SCI. SCI04 had worse proprioceptive sense than SCI 01. For the power traces, positive values represent generative and negative values represent absorptive power. Flexor torques are positive. Arrows indicate regions of interest highlighting the effects of greater proprioceptive deficits on controlling lower limb dynamics. B. Scatter plot of the knee muscle torque produced by each individual with SCI when pointing to the HKA target in relation their movement detection sense score, irrespective of visual condition. The regression lines and R^2^ value were estimated using a linear mixed-effect model fitted to assess the crude relationship between knee muscle torque and movement detection sense. The black dot and error bars represent the mean and standard deviation (SD) of the control data.

## Discussion

In this study, we present a novel lower limb pointing task that manipulated the degree of difficulty by varying the interjoint coordination, and therefore the intersegmental dynamics, required for each target. To control for varying levels of motor impairment among individuals with SCI, we scaled each target to motor function. Our findings show that this task can distinguish motor strategies between individuals with SCI and healthy controls while end-point performance between groups is indistinguishable. Individuals with SCI seem to have difficulties in controlling active and passive knee torques. Through this task, we also identified that these altered motor strategies in those with SCI are associated with greater proprioceptive impairments and end-point performance. Thus, this functional test could be a useful tool for understanding motor strategies and the impact of proprioceptive deficits in people with SCI.

We expected that only the ‘complex’ HKA target would challenge the control of intersegmental dynamics because of the multiple simultaneous joint motions. However, our data indicate that the ‘simpler’ KA or HA target also reveals challenges to intersegmental control. Moving to these targets requires stabilization of one joint while moving the other. Differences between controls and the SCI group were not present at the joint that was the primary mover for the simple targets. Rather, we observed differences at the joint that was to remain stationary. To keep a segment stationary while the adjacent one moves depends on muscle activity produced to control for the passive torques arising from the motion of the moving segment [43]. Indeed, for the HA target, our data suggest that individuals with SCI produce less of the knee extensor muscle torque that would normally control for the passive flexor torque at the knee exerted by gravity. Similarly, the motor strategy for moving to the KA target should involve generating hip extensor muscle torque to maintain the thigh segment in a neutral, vertical position. However, the SCI group had difficulties maintaining sufficient hip extensor muscle torque when moving to the KA target, and some participants even generated hip flexor muscle torque. Thus, the simple targets may help pinpoint whether individuals have difficulties in specifically controlling for the motion of the thigh and/or shank segment. As we discuss below, the complex targets may help identify the compensatory motor strategies adopted by individuals with SCI and those with more severe proprioceptive impairments, as multiple motor strategies can be used to point to this target.

Despite the differences in motor strategies, we found only minimal between-groups differences in end-point performance. This finding is reminiscent of tasks where motor strategies are different but kinematic patterns remain the same. For example, following locomotor training, participants with motor-incomplete SCI exhibit recovery to normative patterns of foot trajectory motion, but not muscle activation patterns, during treadmill walking [44,45]. Muscle activation patterns during gait revealed EMG waveforms that were not only different than controls but also variable across individuals [44,45]. In our data, these variations in motor strategies becomes more apparent upon assessing the spectrum of proprioceptive impairments in individuals with SCI. Specifically, compared to those with milder proprioceptive deficits, individuals with more severe impairments employed distinct motor strategies and these were linked to end-point performance. Thus, the conservation of foot kinematics in walking may reflect a systems-level goal aimed at producing the highly stereotyped kinematic gait pattern [46] and not necessarily applicable to performance of individual participants in the discrete motor tasks utilized here.

We compared task performance with and without vision, expecting that pointing in the obstructed-vision condition, especially to the complex target, would reveal greater deficits in end-point performance compared to full vision. However, we did not find an effect of visual condition even in those with severe proprioceptive impairments. There are at least two potential reasons. First, we gave visual feedback of limb position before movement initiation. Second, the target was always present and remained in sight for the entirety of the movement in both visual conditions. Evidence from previous studies of individuals with large-fiber neuropathy show that even when given only brief visual feedback of their limb at movement onset, they perform similarly to when they are provided limb feedback throughout the movement [15,16]. Thus, the timing with which we delivered visual feedback relative to the onset of movement may have confounded the effects of vision and proprioception in this task. [15,16,47]

Although we could not reveal the effect of proprioceptive deficits by obstructing vision, the heterogeneity in the level of proprioceptive impairment among our participants with SCI helps reveal an association between proprioceptive deficits and compensatory motor strategies.

Proprioceptive sense was associated with muscle torque generation for the HKA and KA targets. When pointing to the KA target, our data indicated a pattern where individuals with worse proprioceptive sense seem to attempt a hip flexor muscle torque strategy, as opposed to maintaining hip extensor muscle torque to stabilize the thigh segment while the knee flexes. This disordered hip flexor muscle torque strategy was associated with greater anterior end-point error and foot-path area. For the HKA target, we found that worse proprioception was associated with less knee flexor muscle torque. Producing less knee flexor muscle torque would attenuate the interaction torques arising from shank angular acceleration and was associated with greater anterior end-point error. Although the loss of muscle force after SCI can also contribute to changes in motor strategy [48], in this study we scaled the targets to each individual’s motor capabilities, suggesting that the compensatory strategies arose from challenges in controlling for intersegmental dynamics. The strategy of limiting knee muscle torque for lower limb movements is reminiscent of findings from individuals with large-fiber neuropathy where gait analysis showed a stiff-knee gait strategy to reduce the degrees of freedom [49]. Indeed, stiff-knee gait patterns are characterized by knee extensor moments during late stance and reduced knee flexor velocity and range of motion during swing [50,51]. During naturalistic reaching, when the need to control intersegmental dynamics increases (i.e., faster reaching), individuals with large-fiber neuropathy also reorganize their motor-control strategy to limit elbow motion [24]. Our task may provide a simple approach for detecting such simplification of motor strategies (i.e., limiting knee motion) in individuals recovering from neurological injury resulting in varying degrees of proprioceptive deficits.

The finding that muscle torque seemed to only associate with MDS, but not JPS, provides preliminary evidence suggesting that JPS and MDS may affect movement control differently. Movement detection sense may associate with motor strategies because muscle activation patterns are responsive to the velocity and acceleration of the different segments (i.e., intersegmental dynamics and interaction torques) [13,25,31]. Investigations in neurologically impaired populations have provided mixed results about the role of JPS and MDS in controlling movement. Studies in adults [52] and children [53] who have suffered a stroke showed that JPS does not associate with indicators of reaching performance. However, studies in adults with SCI show that both JPS and MDS are associated with controlling foot-height in relation to real [19] and virtual [20] obstacles. Further, in older adults, a balance training program improved velocity discrimination at the ankle but not joint position sense, perhaps indicating the importance of velocity-dependent measures of proprioception in controlling movement [54]. Future studies should consider velocity-dependent measures of proprioceptive sense alongside joint position sense when investigating the role of proprioceptive sense depending on the outcome of interest (e.g., accuracy of movements vs. motor control strategies).

Limitations and future considerations of this novel paradigm include task instructions and analysis techniques. First, as discussed above, removing visual feedback of the limb just prior to movement had minimal effect on task performance. To further probe the effect of visual feedback in this task, future studies could manipulate visual feedback of the lower limb for varying time delays before the start of the movement to vary the reliance on proprioceptive feedback to perform this task. Longer delays between obstructing limb feedback and beginning a motor action decreases movement accuracy in individuals lacking proprioceptive sense, presumably from limb drift and the inability to maintain the motor plan [15,16,47]. Second, muscle torque derived from inverse dynamics is a residual term after modeling the effects of gravity and interaction torques at each joint [32]. It therefore reflects a net muscle torque, including active muscle contractions and the effects of other passive structures around the joint, and is unable to distinguish whether the muscle torque was generated from uniarticular or biarticular muscle action. Future studies utilizing EMG and/or biomechanical modelling that includes muscle simulations could help distinguish the relative contributions of biarticular muscles. This will provide more detailed analysis of force distribution across multiple joints, joint coordination, and energy transfer and efficiency [55,56], as well as understanding the role of biarticular muscles in controlling motion-related feedback [57–59]. EMG could further be used to understand the compensatory motor-strategies used by individuals with SCI, such as muscle co-activation.

### Conclusions

We developed a novel lower limb pointing task as a surrogate for assessing the control of intersegmental dynamics during skilled leg movements. This task can distinguish between the motor strategies used by controls and individuals with SCI when pointing to the different targets. Our findings show that pointing not only to complex but also simple targets can challenge intersegmental control and reveal the impact of lower limb proprioceptive deficits on lower limb motor strategies in individuals with SCI. Complex targets help reveal compensatory strategies, whereas simple targets can specifically identify challenges with controlling the motion of the thigh or shank segment. Further development of this paradigm may involve refinement of how and when visual input is obstructed to better discern the global effects of proprioception on task performance. This task can be extended to other neurologically or physically impaired populations to help discern the sensorimotor mechanisms underlying their motor impairments.

## List of abbreviations

10-MWT: 10-meter walk test
AFO: Ankle-foot orthosis
AIC: Akaike Information Criteria
ASIA: American spinal cord injury association
AIS: ASIA impairment scale
FV: Full vision
HA: Hip-ankle
HKA: Hip-knee-ankle
JPS: Joint position sense
KA: Knee-ankle
LEMS: Lower extremity motor score
LMEs: Linear mixed-effects models
MDS: Movement detection sense
N-T: Non-traumatic
OV: Obstructed vision
QQ: Quantile-quantile
SCI-FAP: Spinal Cord Injury Functional Ambulation Profile
T: Traumatic

## Declarations

## Acknowledgments

We thank Maya-Sato-Klemm, Emily Deegan, and Alison Williams for their contributions to data collection. We also thank our participants; this work would not be possible without their time and commitment.

## Author Contributions

RNM, DSM, and TL conceptualized and designed the research. RNM, GE, and MC performed the experiments. RNM and MC conducted the primary analysis. RNM prepared the figures. RNM, DSM, and TL interpreted the results, and drafted and revised the manuscript. All authors contributed to the article and approved the submitted version.

## Funding

We are grateful for the support of the Canadian Institutes of Health Research (CIHR MOP-136914), and the UBC four-year fellowship for PhD students (#6456)

## Availability of data and materials

All data and materials pertaining to this study can be made available upon reasonable request to the corresponding author.

## Ethics approval and consent to participate

All procedures were approved by the University of British Columbia Clinical Research Ethics Board, and all participants provided written informed consent

## Consent for publication

All participants provided written informed consent for the data to be published.

## Competing Interest

The authors declare that they have no competing interests

## Notes

### Competing Interest Statement

The authors have declared no competing interest.

